# Genetic screen to test microRNA function in peripheral glia morphology

**DOI:** 10.1101/2024.04.12.589271

**Authors:** Sravya Paluri, Vanessa J. Auld

## Abstract

Glial cells perform many functions in the nervous system, including maintaining the blood-brain/nerve barriers and structurally supporting axons. While their functions are well-characterized, the complex molecular mechanisms important for their development are less known. Here, we investigated whether microRNA-mediated post-transcriptional regulation is involved during glial development, ensheathment and blood-nerve-barrier formation in *Drosophila*. In this study, we systematically knocked down 120 different microRNAs by competitive inhibition using microRNA-sponges and analyzed peripheral glial morphology. Knockdown of miRNA-125 in the blood-nerve barrier-forming glia (subperineurial glia) resulted in the most penetrant morphological defects. Since microRNA-125 is co-transcribed with miRNAs-let7 and −100 in a genetic cluster, our further verification for subperineurial glia function included miRNA-125 plus all other members of this cluster. However, the loss of each individual gene and the entire cluster did not lead to any morphological defects in the subperineurial glia. To test the efficiency of the microRNA sponge approach in subperineurial glia, we expressed a sponge targeting a microRNA established to be vital for blood-brain barrier formation (microRNA-285) and found no defects in brain lobes and peripheral nerves. Given that a scrambled-sponge control also generated morphological defects, this suggests that using miRNA sponge lines may not be an effective approach to study miRNA function in Drosophila peripheral glia.

## Introduction

In the nervous system, glia perform diverse functions such as providing structural support to axons, protecting the brain from infection and maintaining the blood-brain/nerve barrier (Freeman & Doherty, 2006; Kettenmann & Ransom, 2013; Stork et al., 2012). The blood-nerve barrier (BNB) is formed by tight junctions (TJ) between endothelial cells in vertebrates and pleated septate junctions (pSJ) between glial cells in invertebrates to restrict the paracellular diffusion (Hortsch & Margolis, 2003; Tsukita et al., 2001). However, the mechanisms controlling BNB formation and stability are largely unexplored.

The *Drosophila* peripheral nerve serves as an excellent model for understanding glial function in nerve development and stability. Each nerve consists of three glial layers, which share functional and molecular similarities to vertebrate peripheral glia. The exterior perineurial glia interact with the extracellular matrix (ECM) similar to vertebrates (Freeman & Doherty, 2006; Olsson, 1990; Xie & Auld, 2011). The subperineurial glia (SPG) in the middle layer form the blood-nerve-barrier (BNB) through pSJs, which are molecularly conserved with the paranodal junctions found in vertebrate myelinating Schwann cells (Bhat, 2003). The core *Drosophila* SJ consists of NeurexinIV, Coracle, Neuroglian and Contactin, which are conserved with paranodal junction components Contactin-associated protein (Caspr), Band 4.1, Neurofascins, and Contactin respectively (Banerjee et al., 2006.; Bellen et al., 1998; Bhat, 2003; Boyle et al., 2001; Charles et al., 2002; Faivre-Sarrailh et al., 2000; Tait et al., 2000). Loss of any core SJ proteins compromises the pSJ integrity leading to loss of the permeability barriers including the BNB and lethality (Auld et al., 1995; Baumgartner & Troy, 1996; Fehon, 1994). The SPG ensheathes the innermost layer of glia formed by the wrapping glia (WG), which wrap around bundles of peripheral axons similar to the vertebrate non-myelinating Schwann cells that form ‘Remak’ bundles (Stork et al., 2008, Nave, 2010). Loss of WG ensheathment of axons alters the animal locomotion (Kottmeier et al., 2020; Matzat et al., 2015), while their ablation leads to lethality (Petley-Ragan et al., 2016). Therefore, proper development and function of all the peripheral glia layers is crucial for animal survival. While the basic process of glial development is known, the regulatory mechanisms involved in coordinating cohorts of mRNA and protein expression during development are not.

Spatio-temporal expression of the appropriate cohorts or clusters of genes is essential for proper development and differentiation (Dong & Liu, 2017). One mechanism to regulate a set of proteins is through microRNAs (miRNAs). miRNAs regulate mRNA transcripts by binding to a highly specific sequence in the 3’UTR region to either trigger mRNA degradation or translational repression (Ha & Kim, 2014). In vertebrates, select miRNAs control the development and differentiation of glia. For example, microRNA-23a regulates the expression of proteins crucial for oligodendrocyte glia (OL) differentiation and myelination (Lin et al., 2013). Ablation of the microRNA processing protein, Dicer, post-developmentally in OL results in a neurodegenerative disease accompanied by demyelination in the brain (Shin et al., 2009). Similarly, in the peripheral nervous system (PNS), deletion of Dicer in premature Schwann cells leads to impairment in differentiation and myelination (Bremer et al., 2010; Pereira et al., 2010). This loss of Dicer also leads to axonal degeneration, suggesting that Schwann cell-axon interactions also require miRNAs (Pereira et al., 2010). While miRNAs are known to control some vertebrate glial development, their role in the formation of Schwann cell junctions and in non-myelinating Schwann development and maintenance is not known. To address these questions, we used *Drosophila* as a model to identify and investigate the roles of miRNAs in peripheral glial development.

In *Drosophila* different microRNA regulatory networks affect processes like lifespan, locomotion, homeostasis, circadian rhythm, neurogenesis and neuromuscular junction development (Foo et al., 2017; Loya et al., 2014; Mazaud et al., 2019; Tsai et al., 2019; Tsurudome et al., 2010; You et al., 2018; Yuva-Aydemir et al., 2015). In the central nervous system (CNS), miRNA-285 was identified as a key regulator of the Hippo signaling pathway necessary for SPG ploidy and glial growth and expansion (Li et al., 2017). However, in the PNS, the identity and function of miRNAs necessary for SPG and WG development are unknown. Therefore, we employed a genetic loss-of-function method to individually downregulate multiple microRNAs in a genetic screen and tested for changes to SPG and WG morphology.

## Materials and methods

### Drosophila strains

The following lines were obtained from the Bloomington stock center (BDSC): *UAS-mCD8::RFP* and the 120 microRNA-Sponge lines including *miRNA-scrambled-SP* lines (Fulga et al., 2015). Other lines used included: *UAS-miR-9a-SP* (Fulga et al., 2015), *Gli-Gal4* (Sepp & Auld, 1999)*, SPG-GAL4* (Schwabe et al., 2005), *NrxIV::GFP* (Buszczak et al., 2007), *Nrv2-Gal4* (Sun et al., 1999), *Nrv2::GFP* (Morin et al., 2001) and *UAS-mCD8::GFP* (Lee & Luo, 1999), *ΔmiR-let7, ΔmiR-100, ΔmiR-125, Δlet7cKO2* and *Δlet7cGK1* (Sokol et al., 2008). The *UAS-miR-125::dsRed* and *UAS-miR-100::dsRed* lines (Schertel et al., 2012) were ordered from Zurich ORFeome Project (FlyORF).

### Immunolabelling and microscopy

*Drosophila* strains were raised at 25°C unless noted. For the sponge screen, third instar larvae were dissected and were fixed in 4% paraformaldehyde in Phosphate Buffer Saline (PBS) for 20 minutes and subsequently washed with PBS three times. Larvae were then equilibrated first in 70% glycerol and then in Vectashield (Vector laboratories). For the immunostaining protocols, a previously established protocol was used (Sepp et al., 2000). Primary antibodies used included: rabbit anti-Nervana 2.1 (1:1000; Sigma Millipore) and mouse anti-Futsch/22C10 (1:2000; Developmental Studies Hybridoma Bank). Secondary antibodies included goat anti-rabbit conjugated with Alexa 564 and goat anti-mouse conjugated with Alexa 647 (Molecular Probes). After mounting, fluorescent images were taken using a 60x oil immersion objective (NA 1.4) on the Deltavision microscope (Applied Precision/GE Healthcare, Mississauga, Ontario, Canada) with 0.2 μm slices. Deconvolution using SoftWorx was carried out using a point-spread function measured with a 0.2 μm bead in Vectashield (Vector Laboratories). Each image represents a single Z slice and image processing was carried out using ImageJ/Fiji software (Schneider et al., 2012) and figures were compiled using Adobe Photoshop (Adobe Systems).

### Genetic screen phenotype classification and Statistical data analysis

For each of the 120 sponge lines, 5 or more larvae were analyzed. To compare miR-125SP line to the control, the number of larvae (N) showing each category of phenotype was plotted using Prism 9.0 software. Statistical analysis was performed using two-tailed Mann-Whitney U test on Prism 9.0 software (GraphPad Software Inc.) to compare the total percentage of nerves affected for each larva compared to the control.

## Results

### A genetic knockdown screen of individual miRNAs in subperineurial glia

To determine the role of miRNAs in peripheral glial development, we systematically blocked different miRNAs (Table 1) in the subperineurial glia (SPG) by expressing miRNA sponges (*UAS-miR-SPs*) (Fulga et al., 2015). Each miRNA sponge contains sequences complementary to miRNA seed sequences (region that binds to the target mRNA), such that sponge expression leads to the binding and sequestering of that miRNA (Loya et al., 2009). This results in competitive inhibition of miRNAs binding to target mRNAs, resembling loss of function mutants (Fulga et al., 2015). During the screen, 120 different sponge lines were expressed in the SPG using *Gliotactin-GAL4 (Gli-Gal4*) along with a red, or green fluorescent membrane marker (*UAS-mCD8::RFP; UAS-mCD8::GFP*) to assess defects in SPG morphology. Any morphological changes to the wrapping glia (WG) were assessed using Nervana2 endogenously tagged with GFP (*Nrv2::GFP*). Morphological changes were assessed at the wandering third-instar larval stage. Controls were the driver line cross to a scrambled sponge (*Gli-Gal4>ScramSP*), which contains a sequence not specific for any known miRNA.

**Table 1.**
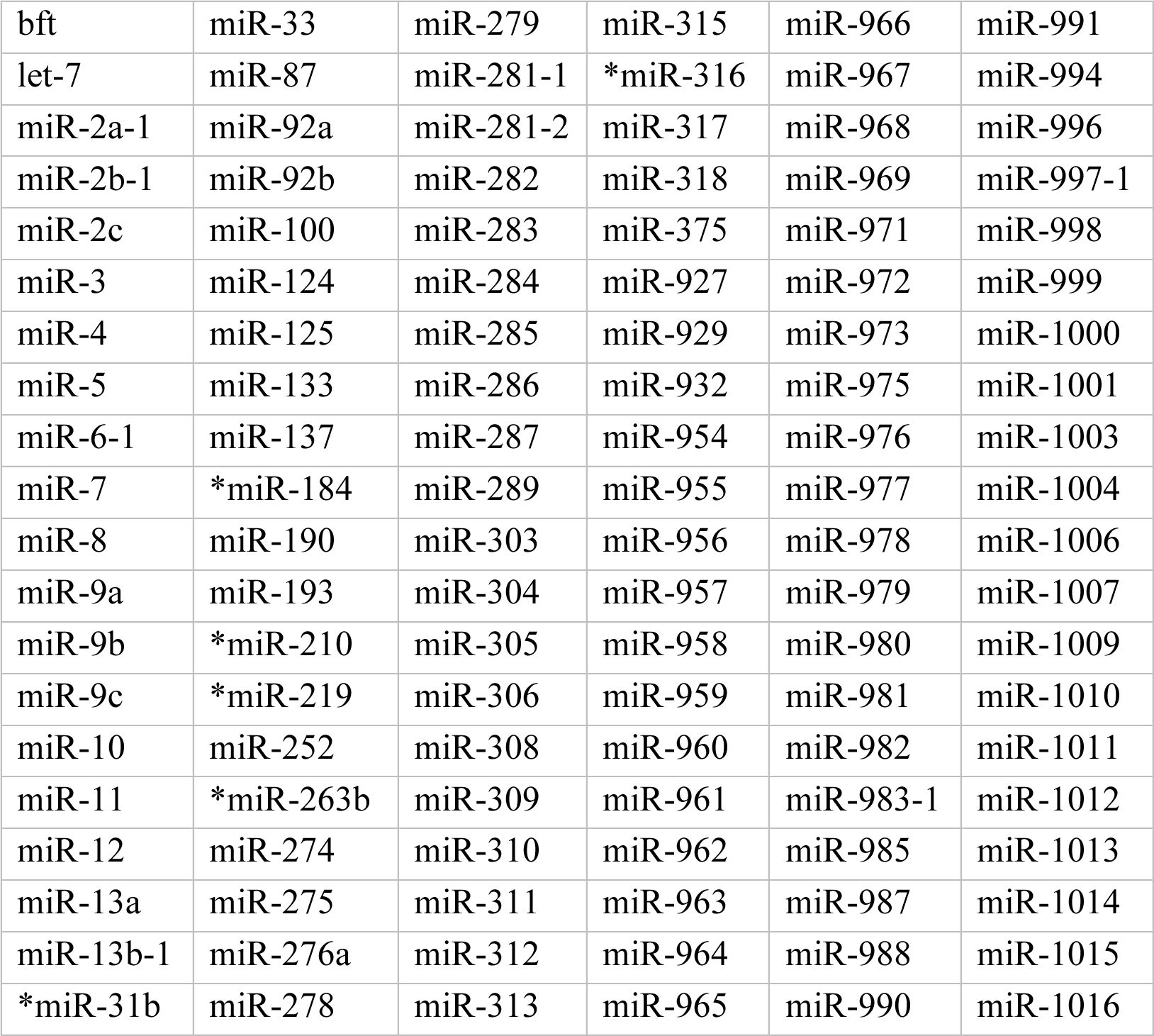
List of microRNAs tested in the screen. miRNAs were competitively inhibited by UAS-miR-SPs driven in SPG using the SPG-specific driver, *Gli-GAL4* for these 120 transgenic lines. SPG were visualized using the fluorescent membrane marker (*UAS-mCD8::RFP*) and WG visualized using Nervana2 endogenously tagged with GFP (*Nrv2::GFP*). Control: The driver line (*Gli-Gal4>UAS-mCD8::RFP, Nrv2::GFP*) crossed to *UAS-Scram-SP*. Experimental lines: the same driver line crossed to the respective sponge lines (*UAS-miR-SPs*). *marked miR-SPs were tested using the same SPG driver line with a different fluorescent marker (*mCD8::GFP*) crossed to \**UAS-miR-SPs* in experimental, and to *UAS-Scram-SP* in control. No. of larvae tested for all crosses > 5.

In the control (*Gli-Gal4>mCD8::RFP; Gli-Gal4>mCD8::GFP*), the SPG layer forms a thin glial layer (white arrowhead) surrounded the underlying WG (yellow arrowhead), which ensheath bundles of peripheral axons (Fig 1). Out of the 120 miRNA sponges tested in our initial screen, 119 sponge lines had no phenotypes and were characterized by normal SPG and WG membrane morphology. However, expression of the miR-125 sponge (*UAS-miR-125SP*) yielded a consistent and higher penetrance of phenotypes in both SPG and WG (Fig 1B-C), with 92% of larvae (24 larvae) displaying a phenotype. The SPG and WG defects (white arrows) included discontinuous, disintegrated, and swollen membranes of both glial layers (Fig 1B, E-J). The defects exclusive to WG (yellow arrowheads) included vacuole formation, aggregation, less WG strands and processes (WG strand extensions) formation (Fig 1H-J). However, in none of the larvae did we find phenotypes exclusive to the SPG layer. The percentage of nerves affected in individual larvae ranged from as low as 6% to as high as 91% with an average of 30% nerves affected per larva (Fig 1C) and these phenotypes were significantly higher compared to control. Surprisingly with the expression of the control scrambled sponge (*Gli-Gal4>UAS-ScramSP*), 47% of larvae displayed some degree of phenotype in either the SPG or WG (32 larvae) (Fig 1B-C). The percentages of nerves affected in individual larvae ranged from 8% to as high as 71% with an average of 15% nerves affected per larva (Fig 1C). Of note, our observations are consistent with previous studies where expression of the scrambled sponge in glial cells results in observable phenotypes (You et al., 2018).

**Figure 1.**
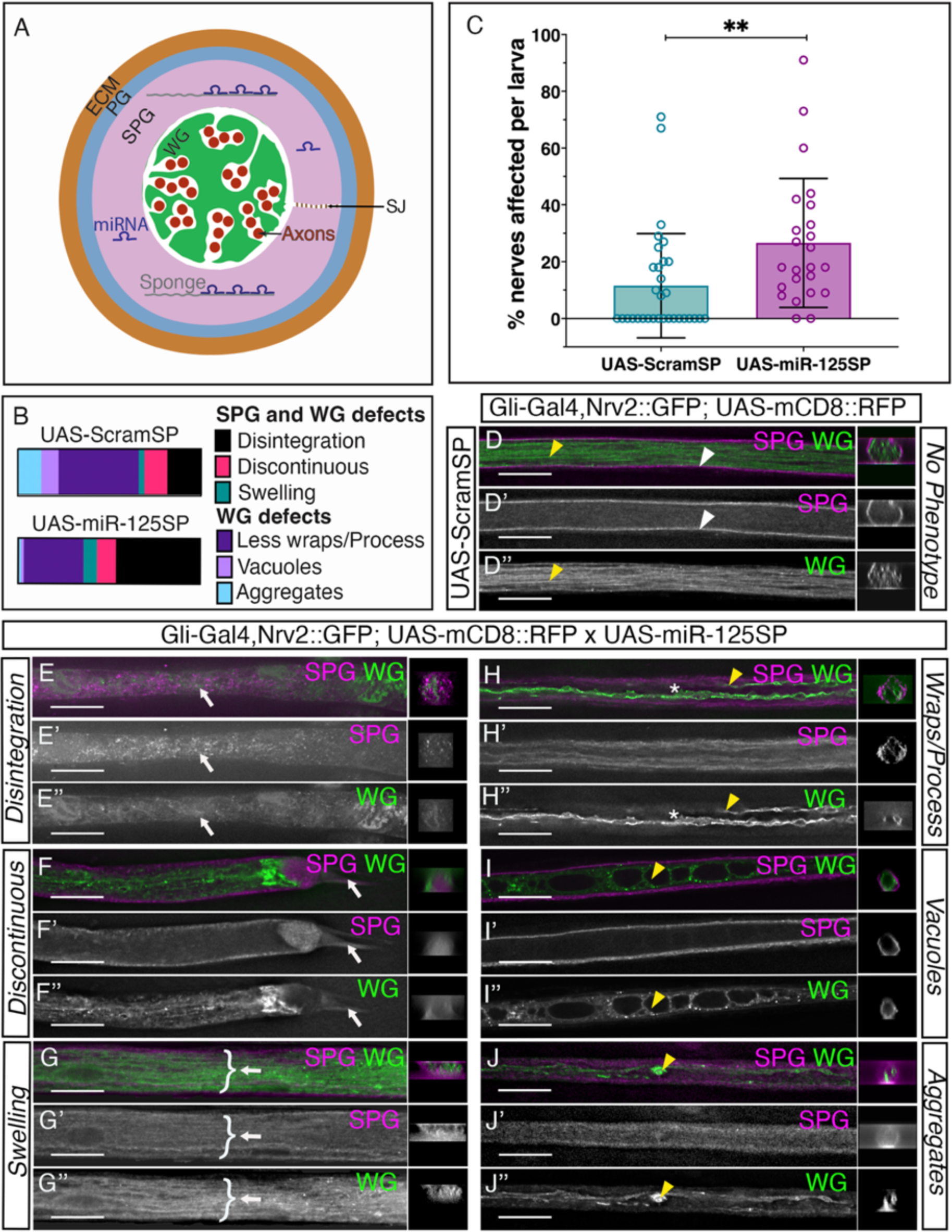
Range of phenotypes observed during the screen. **(A)** Model depicting our genetic screen. *Drosophila* peripheral nerve of third instar larvae with perineurial (PG, blue), subperineurial (SPG, pink) and wrapping glia (WG, green) layer. PG interact with the extracellular matrix (brown), SPG form septate junctions (SJ) and WG wrap axon bundles (red). miRNA sponges (grey) were expressed to competitively inhibit different miRNAs (dark blue) in SPG. **(B-C)** Out of the 120 miRNA sponges tested, miR-125SP resulted in the most penetrant phenotypes compared the control. **(B)** The ratio of phenotypes observed in *UAS-miR-125SP* and the control *UAS-ScramSP* nerves. Defects observed in both SPG and WG: disintegration, discontinuity, swelling. Defects observed in the WG only: reduced wraps/processes, vacuoles, and aggregates. **(C)** Quantification of percentage (%) of affected nerves per larva for the control Scram-SP (32 larvae) and the miR-125SP (24 larvae). Error bars show the standard deviation from the mean. Statistical significance determined by Mann-Whitney U test, P value=0.001. **(D-J)** The SPG driver *Gli-Gal4* driving the expression of *mCD8::RFP* (white arrowheads, magenta) with the WG labelled with *Nrv2::GFP* (yellow arrowheads, green). **(D)** Control: *UAS-Scram-SP*: Representative image of a nerve with no phenotype showing the thin magenta SPG membrane (white arrowhead) surrounding the green WG strands (yellow arrowhead) that ensheathes peripheral axons. **(E-J)** Range of phenotypes observed upon *UAS-miR-125SP* expression in both SPG and WG (white arrows): **(E’, E’’)** Disintegration, **(F’,F’’)** discontinuity and **(G’,G’’)** swelling of SPG and WG membrane, and WG exclusively (yellow arrowheads): **(H’’)** less WG strands and processes formation, **(I’’)** vacuole formation and **(J’’)** WG membrane aggregation. A cross-section of the corresponding transverse nerve is shown on the right. Scale bar=15 μm.

Upon comparing the phenotypes observed between the miR-125 sponge and the scrambled sponge, there were qualitative differences (Fig 1B). With the scrambled sponge, the phenotypes observed were predominantly in the WG, including loss of WG strands. However, with the miR-125 sponge, SPG and WG disintegration was the most prevalent phenotype. Regardless, the expression of the miR-125 sponge in SPG did not affect adult viability. This finding was consistent with another study that showed that ubiquitous expression of this sponge did not affect adult viability (Fulga et al., 2015). Therefore, even though the miR-125 sponge had strong effects on SPG morphology in 30% of the nerves on average, it did not lead to lethality suggesting that the larvae are able to compensate for disruption of the glia.

Overall, our genetic screen revealed that expression of the miR-125 sponge in SPG leads to phenotypes in peripheral glia. Additionally, we found that expression of the control scrambled sponge also resulted in a mild degree of phenotypes, which was still statistically significantly lower than the miR-125 sponge. Therefore, we continued investigating the role of miR-125 in peripheral glia.

### Downregulation of miR-125 in WG does not affect ensheathment of axons

We next investigated if miR-125 is required in the WG. We expressed the miR-125 sponge using a WG-specific driver, *Nrv2-Gal4*, and visualized the membranes using a GFP-tagged membrane marker (*mCD8::GFP*) (Fig 2). In the control (*Nrv2-Gal4>mCD8::GFP, UAS-ScramSP*), WG strands were visualized as multiple thin fine membranes that run parallel to the nerve (white arrowheads) with the expression of Scrambled-sponge visualized using a mCherry tag (Fig 2A). We found that the expression of the miR-125 sponge sequence (*Nrv2>mCD8::GFP, miR-125SP*), along the WG (white arrowheads) had no effect on WG membrane integrity (Fig 2B). Thus, downregulation of miR-125 in WG had no effect on the WG ensheathment of axons.

**Figure 2.**
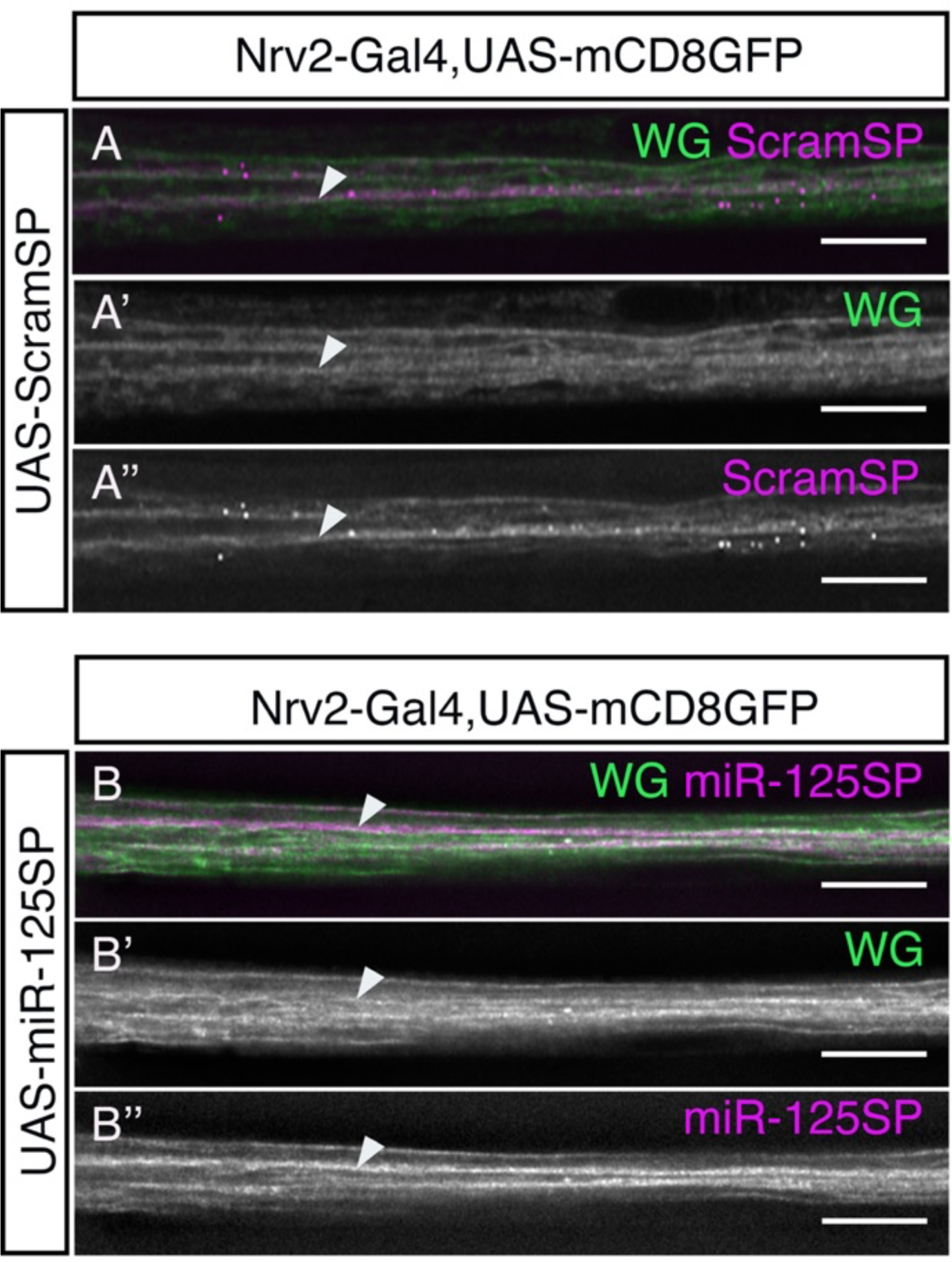
miR-125 sponge expression in the WG has no effect on WG morphology. *Nrv2-Gal4>UAS-mCD8::GFP* (green) was used to visualize the WG morphology (arrowheads), and each miRNA-SP transgene is visualized with mCherry (magenta). **(A-A’’)** Control: *Nrv2-Gal4>UAS-ScramSP* shows no (A’) WG morphology defects with (A’’) ScramSP expression (53 nerves in 5 larvae). **(B-B’’)** Experimental: *Nrv2-Gal4>UAS-miR-125SP* did not generate (B’) WG morphological defects with (B’’) miR-125SP expression (107 nerves in 9 larvae). Scale bar= 15 μm

### miR-125 loss of function mutants do not show SJ morphological defects

Previous studies showed that miR-125 is enriched in a range of vertebrate glia including oligodendrocytes, astrocytes, Müller glia (Lecca et al., 2016; Pogue et al., 2010; Shenoy et al., 2015; Wohl & Reh, 2016) and in neurons (Akerblom et al., 2014; Le et al., 2009) and thus, we continued investigating miR-125 for its potential role in glial cell development and function. Several studies have failed to phenocopy results obtained from sponge expression with their corresponding null mutant (Fulga et al., 2015). Therefore, our primary step was to check if we could phenocopy the miR-125 sponge phenotype using null mutants. microRNA-125, microRNA-let7 (miR-let7) and microRNA-100 (miR-100) are co-transcribed from the same genomic locus, the *lethal-7-cluster* (*let-7-c)* (Chawla et al., 2016; Prochnik et al., 2007; Sempere et al., 2003; Sokol et al., 2008). Our initial miR-sponge screen had found that expression of the miR-let7 and miR-100 sponges had no effect on SPG morphology (white arrowhead, Fig 3). However, because let-7-c miRNAs can cross-regulate each other (Chawla et al., 2016)., we investigated the role of the entire cluster in peripheral glia development. We used two independently generated knockout lines, *Δlet7cKO2* (Chawla et al., 2016) and *Δlet7cGK1* (Sokol et al., 2008). As well we used transheterozygous knockout mutant *Δlet7cKO2/GK1* flies containing rescue transgenes of the other two miRNAs to generate *ΔmiR-let7, ΔmiR-100, ΔmiR-125* respectively (Chawla et al., 2016).

**Figure 3.**
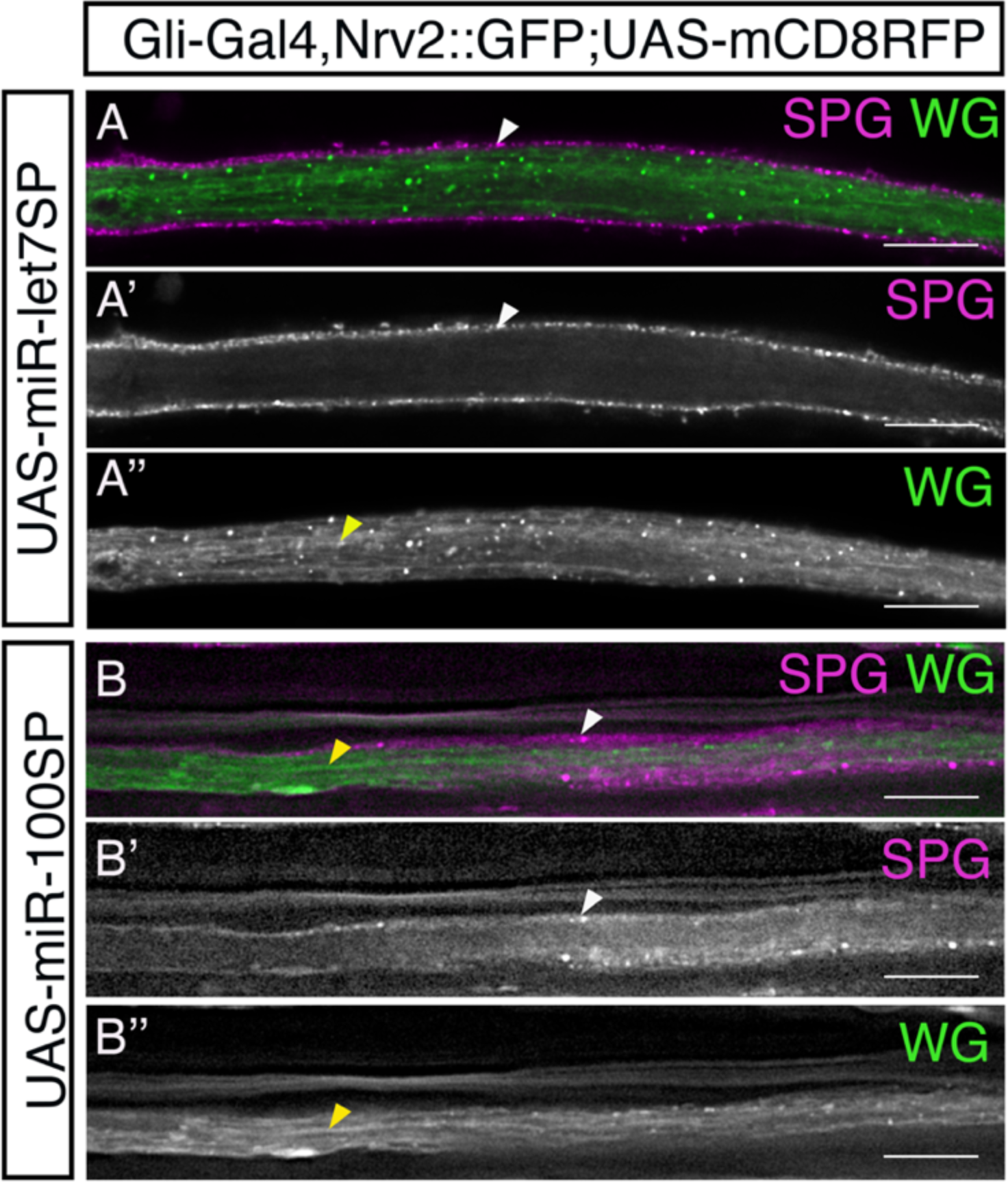
Downregulating miR-let7 and miR-100 by sponge in SPG has no effect on SPG and WG morphology. *Gli-Gal4>UAS-mCD8::RFP* (red) with Nrv::GFP was used to visualize the SPG (white arrows) and WG (yellow arrows) morphology respectively. Downregulation of miR-let7 sponge and miR-100 sponge are compared to Scram-SP control in Fig 1D-D’’. **(A-A’’)** *UAS-miR-let7 sponge* (UAS-miR-let7SP) exhibited no **(A’)** SPG or **(A’’)** WG defects (58 nerves in 6 larvae). **(B-B’’)** *UAS-miR-100 sponge* (UAS-miR-100SP) exhibited no **(B’)** SPG or **(B’’)** WG defects (62 nerves in 7 larvae). Scale bar= 15 μm

To confirm the necessity of the *let-7-c* cluster in the SPG, we assessed pSJ morphology by visualizing the septate junction protein Nervana 2.1 (Nrv2.1) as SPG morphological defects compromise pSJ integrity. Given that the pSJ proteins are highly interdependent (Bätz et al., 2014; Baumgartner et al., 1996; Behr et al., 2003; Genova & Fehon, 2003; Hall et al., 2014; Nelson et al., 2010; Paul et al., 2003; Stork et al., 2008; Wu et al., 2004), the Nrv2 expression pattern serves as an indication of pSJ morphology. We also tested peripheral axonal integrity, using the anti-Futsch (22C10) antibody. In *w^1118^* controls, the pSJs were confined to a thin boundary along the length of each SPG and at the points of contact of two SPG cells (yellow arrowhead), whereas axon bundles were observed as multiple thin straight lines expressed along the nerve (Fig 4N). The SJ and axons of individual *ΔmiR-let7*, *ΔmiR100* and *ΔmiR-125* mutant larvae did not show any defects compared to the control (Fig 4K-M). Due to the potential cross-regulation of members of this cluster (Chawla et al., 2016), we hypothesized that the loss of these individual miRNAs might be compensated by the others. Therefore, we tested the effect of loss of the entire let-7 genomic cluster using two different *let-7-cluster* mutants, *Δlet-7c-GK1* and *Δlet-7c-KO2* (Sokol et al., 2008). Homozygous mutants of the *Δlet-7c-GK1* and *Δlet-7c-KO2* did not exhibit any SJ or axonal defects (Fig 4H, I). In addition, the transheterozygous combination of *Δlet-7c-GK1/ Δlet-7c-KO2* did not have any SJ or axonal defect compared to the control *w^1118^* (Fig 4J). Thus, our studies revealed that loss of the entire *let-7-c* cluster does not affect pSJ or peripheral axonal integrity. The fact that *ΔmiR-125* mutant larvae did not exhibit any pSJ defect formed by SPG contradicted our findings from the sponge screen that showed that reduction of miR-125 affected SPG morphology.

**Figure 4.**
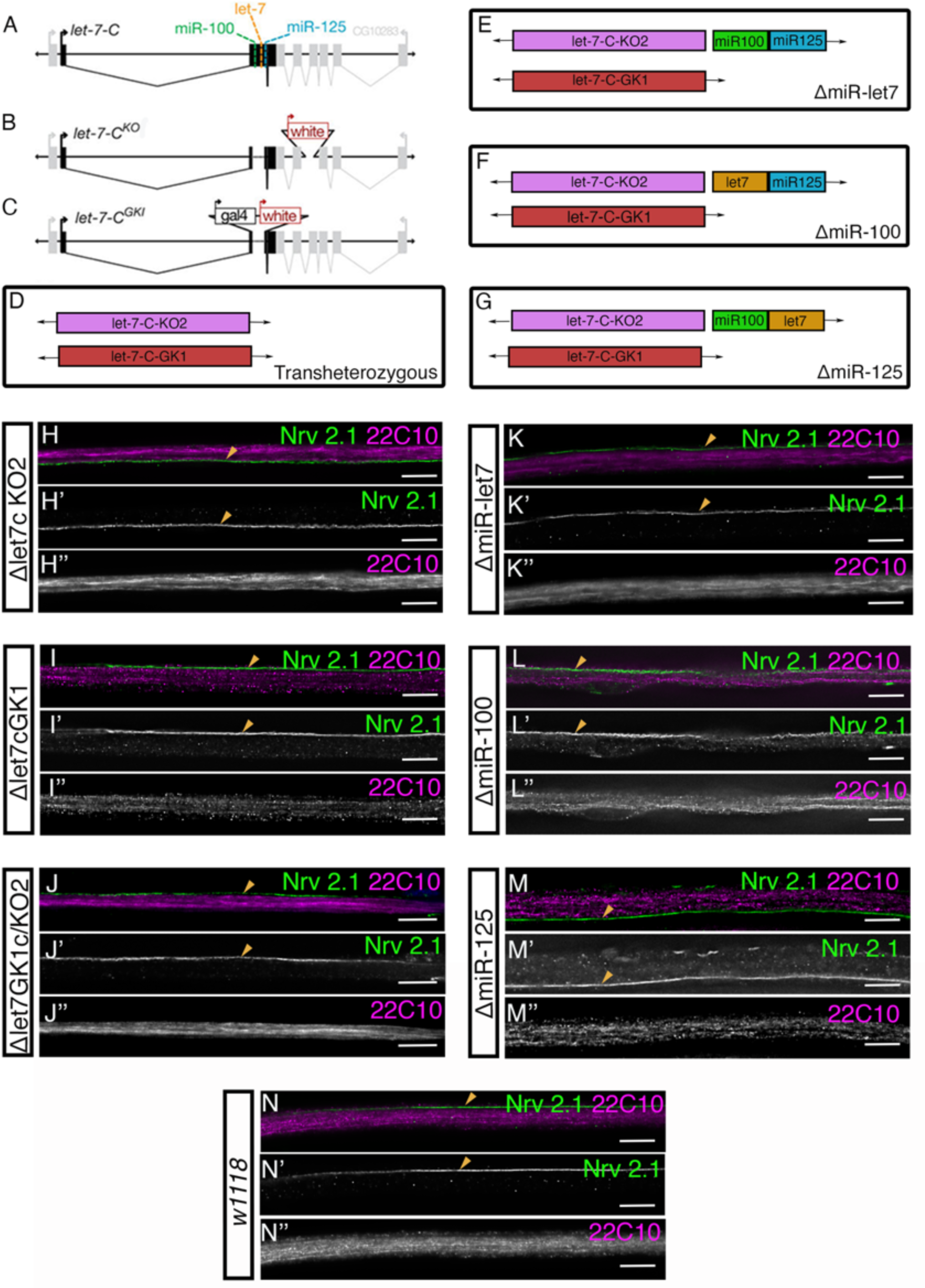
*let-7-c* knockout microRNAs have no septate junction or axonal morphological defects. **(A)** *let-7-c* gene encodes three microRNAs-miR-100, miR-125 and miR-let7 (modified from Sokol et al., 2008). **(B,C)** Two independent complete knockout null mutants generated by ends-out-homologous recombination *Δlet-7-cKO2* and *Δlet-7-c-GK1* (Sokol et al., 2008) were used for generating **(D)** transheterozygous mutant *Δlet-7-GK1c/KO2* and for **(E,F,G)** independent null mutants generated in the transheterozygous mutants by reintroducing two of the other microRNAs. **(E)** *ΔmiR-let7* was reintroduced with miR-100 and miR-125. **(F)** *ΔmiR-100* was reintroduced with miR-let7 and miR-125. **(G)** *ΔmiR125* mutant was reintroduced with miR-let7 and miR-100. **(N)** In the control (*w^1118^*), SJ (Nrv2.1; green) formed a thin line along the length of the SPG (yellow arrow) and the axons (magenta) immunolabeled for Futsch (22C10) formed bundles running parallel to the nerve. **(H-J)** Homozygous mutants with loss of **(H-H’’)** *let7-c, let7cKO2* and **(I-I’’)** *let7GK1c* and **(J-J’’)** both *let7GK1c/KO2* displayed no defects in SJ and axonal morphology. **(K-M)** Individual homozygous null mutants **(K-K’’)** *ΔmiR-let7*, **(L-L’’)** *ΔmiR-100* and **(M-M’’)** *ΔmiR-125* had no defects in SJ and axonal morphology. Scale bar= 15 μm.

### Overexpression of miR-125 and miR-100 does not affect SPG morphology

As a final test of *let-7-c* function in peripheral glia, we then overexpressed miR-100 and miR-125 in SPG to analyze SPG morphology integrity. The *Gli-GAL4* driver expressing a GFP membrane marker (*mCD8::GFP*) was crossed to lines with each miRNA, *UAS-miR-100::dsRed* and *UAS-miR-125::dsRed* (Schertel et al., 2012) or to *w^1118^* as the control. In the control (*Gli-Gal4>mCD8::GFP*), the SPG membrane formed a thin membrane surrounding each nerve (Fig 5C). The presence of the *UAS-miR-125* and *UAS-miR-100* was visualized using dsRed (yellow arrowhead) along the SPG membrane (Fig 5A’’, B’’). We found that overexpression of miR-125 (*Gli-Gal4>mCD8::GFP, miR-125*) and miR-100 (*Gli-Gal4>mCD8::GFP, miR-100*) in SPG had no effect on SPG membrane integrity (Fig 5A, B). Taken together, neither loss nor overexpression of *let-7-c* led to defects in SPG morphology or formation.

**Figure 5.**
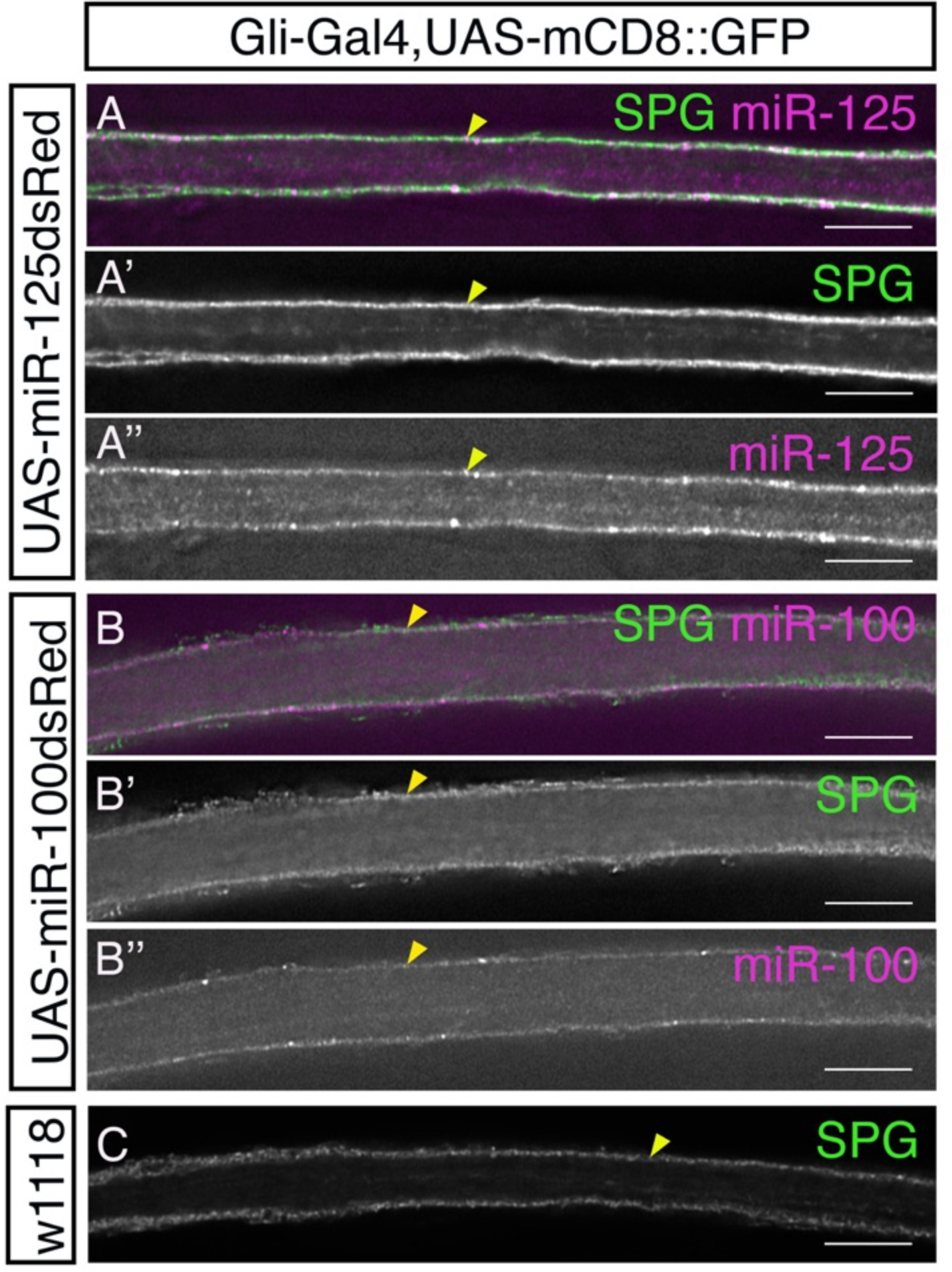
Overexpression of miR-100 and miR-125 had no effect on SPG morphology. *Gli-Gal4>UAS-mCD8::GFP* (green) was used to visualize SPG membrane morphology and drive each miRNA transgene expressing dsRed (magenta). Expression of **(A-A’’)** *UAS-miR-125* (151 nerves in 15 larvae) or **(B-B’’)** *UAS-miR-100* (136 nerves in 10 larvae) did not result in any SPG morphological defects and membranes appeared similar to control **(C)** *w^1118^*(19 nerves in 2 larvae). Scale bar= 15 μm

### let-7-complex expression was not detectable in the peripheral nerve membrane

Having established that miR-125 function is not required in the SPG or WG for their respective morphology and function, we wanted to check if miR-125 or the let-7-c is expressed in peripheral glia. Previous studies revealed that let-7-c is expressed in the adult brain, ventral nerve cord, motoneurons, some abdominal muscles, sensory organs in the head, flight muscles, alimentary canal, and reproductive tracts of male and female flies (Sokol et al., 2008).

For analysis of the expression of *let-7-c* in the peripheral nerve of third-instar larvae, the *Δlet-7-c GK1* mutant line (Sokol et al., 2008) containing a Gal4 sequence was crossed to a fluorescent reporter (*UAS-mCD8::GFP*) and the pSJs were visualized using the Nervana 2.1 (Nrv2.1) antibody (Fig 6). We failed to observe let-7-c-driven GFP expression within the peripheral nerve (white asterisks), which matched prior work that found endogenous *let-7-c* levels are low in the larval CNS (Wu et al., 2012). Overall, the expression of the *let-7c cluster* is either absent or below the levels of detection in the peripheral nerves of third-instar larvae.

**Figure 6.**
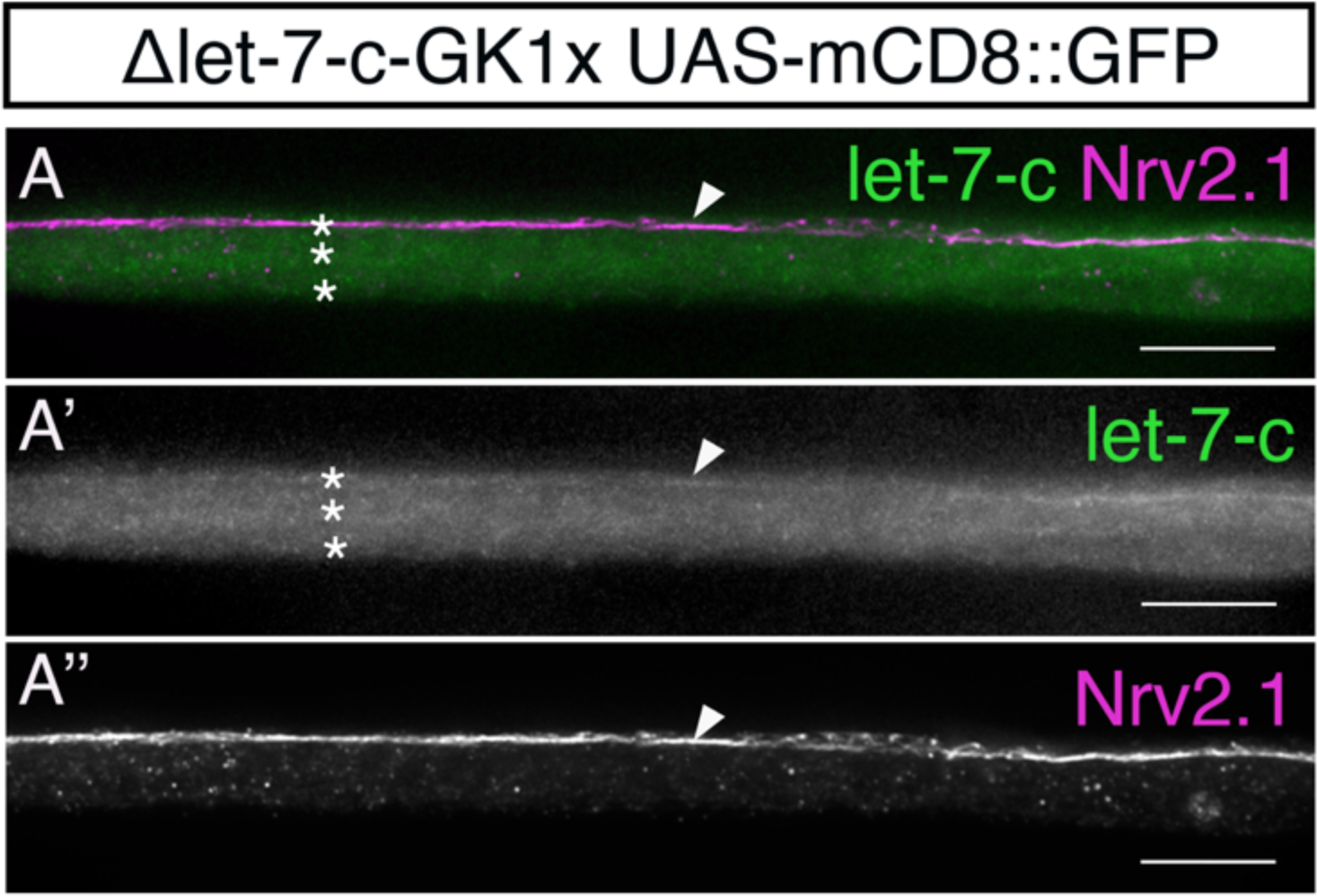
Lethal-7-complex (let-7-c) is not expressed in glial membranes of the peripheral nerve. **(A-A’’)** *Δlet-7-c-GK1* (Sokol et al., 2008) containing a Gal4 sequence was crossed to *UAS-mCD8::GFP* and counterstained with Nrv2.1 to visualize the SJ marker Nrv2. **(A’)** shows let-7-c expression is absent along the peripheral nerve membrane (asterisks) with some background expression in some regions of the SPG membrane (arrowhead). Scale bar= 15 μm

Overall, our mutant analysis revealed that miR-let-7, miR-100 and miR-125 are individually and collectively not required for septate junction or axonal integrity and likely not even present in peripheral glia in third instar larval stages. These results suggest that our sponge screen likely generated false-positive results.

### miR-285 knockdown by sponge does not affect the blood-brain barrier

As the sponge screen appeared to generate false positive results, we sought to test the efficiency of the sponge lines using a miRNA with a known function in SPG. Third-instar larvae lacking miR-285 have a defective CNS blood-brain barrier due to a dysfunctional Hippo signaling pathway (Li et al., 2017). To test the efficiency of the sponges in SPG, we downregulated miR-285 expression by using a miR-285 sponge (*UAS*-*miR-285SP*) (Fulga et al., 2015) and analyzed the pSJ structure (Fig 7). In the scrambled sponge control (*SPG-Gal4>mCD8::RFP, UAS-Scram-SP*), the brain lobes and peripheral nerves had intact pSJs visualized by NrxIV::GFP in the CNS or Nrv2.1 in peripheral nerves (yellow arrowheads, Fig 7A-B). However, we found that in the larvae expressing the miR-285 sponge (*SPG-Gal4>mCD8::RFP, UAS-miR-285SP*), the pSJs were similar to the control in both the brain lobes and peripheral nerves (Fig 7C-D). Therefore, we were unable to phenocopy using the sponge lines the blood-brain barrier defects seen with loss of function mutants from the previous study (Li et al., 2017).

**Figure 7.**
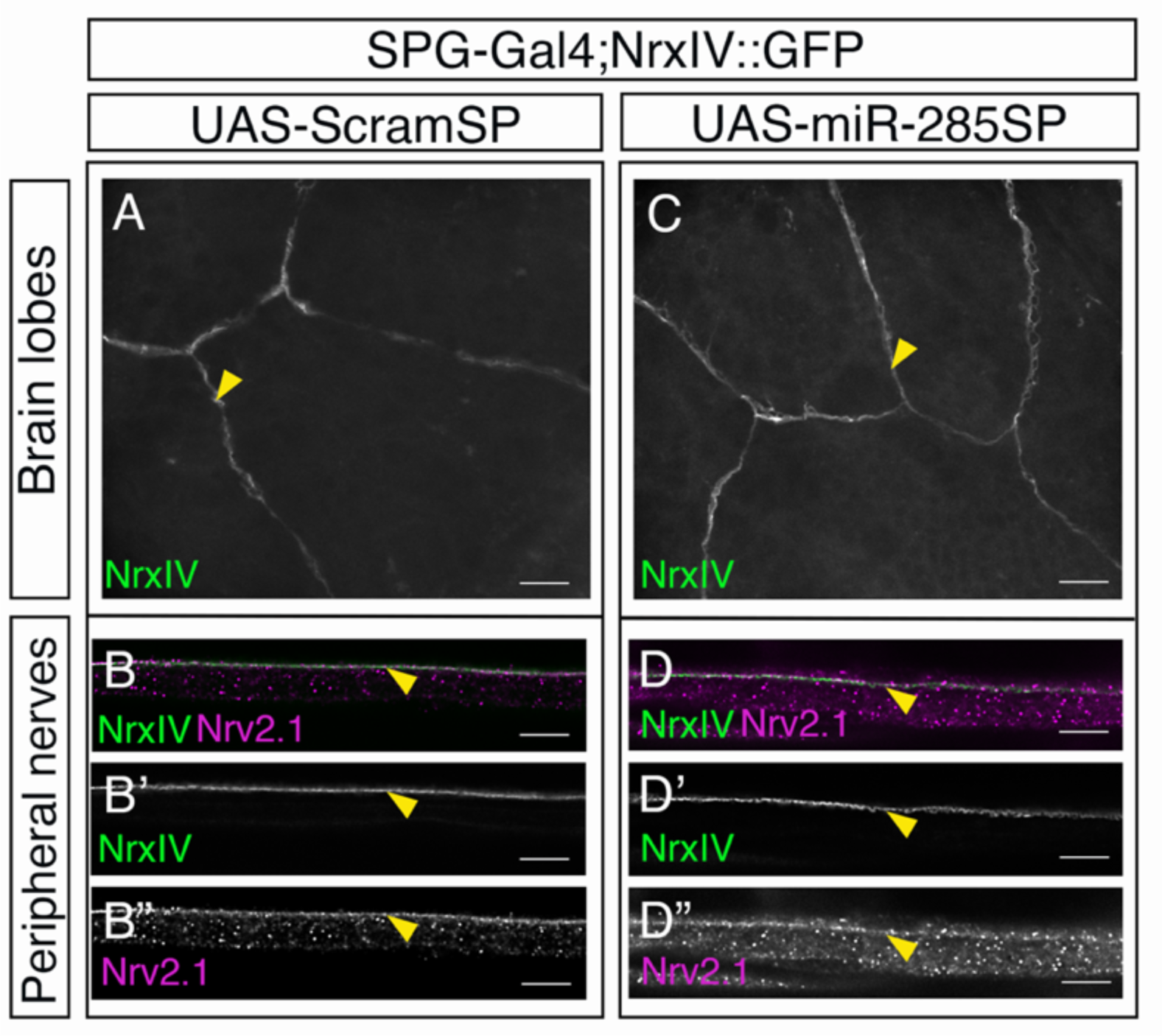
Expression of the miR-285 sponge in SPG had no effect on septate junctions in the brain lobes or in the peripheral nerves. SPG-Gal4 was crossed to the control, scrambled sponge (*UAS-ScramSP*) and to the miR-285-sponge (*UAS-miR-285SP*). Two markers were used to visualize the pSJs, *NrxIV::GFP* (green) in the brain lobes and Nrv2.1 immunolabeling (magenta) in the peripheral nerves. (**A**) Brain lobes of the control *UAS-ScramSP* (4 larvae) with NrxIV expression marking the pSJs (arrowheads). **(B-B’’)** Peripheral nerves of the control *UAS-ScramSP* with NrxIV expression marking the pSJs (arrowheads) and colocalizing with Nrv2.1 (43 nerves in 4 larvae). **(C)** Brain lobes with overexpression of *UAS-miR-285SP* in the SPG (6 larvae) had no pSJ phenotypes when visualized with NrxIV (arrowheads). (**D-D’’**) Peripheral nerves with overexpression of *UAS-miR-285SP* in the SPG had no defects in the pSJ visualized with NrxIV and Nrv2.1 (arrowheads) (82 nerves in 6 larvae). Scale bar= 15 μm

Overall, our results point to the difficulty in using the sponge lines to test for microRNA function in the SPG. Although our study failed to identify which miRNAs are important in peripheral nerve, our study highlighted that the let-7-cluster is not required for pSJs and SPG.

## Discussion

In *Drosophila melanogaster*, about 100 miRNAs play an important role in at least one biological function including survival, locomotion, fertility and maintenance of the permeability barriers (Chen et al., 2014). In the nervous system, miRNAs play an important role in glial homeostasis in the adult brain and protect glia from excitotoxicity (Aw et al., 2017; Foo et al., 2017). However, their role in the formation of peripheral glia has not been characterized.

Overall we found that the downregulation of 119 microRNAs using the sponge collection (Fulga et al., 2015) in SPG did not affect SPG or WG morphology. However, the expression of the miRNA-125 sponge generated consistent and penetrant phenotypes in both the SPG and WG. However, a miR-125 loss of function mutant or loss of the entire let7-c cluster of miRNAs, including miR-let7, −125 and −100 did not affect peripheral pSJ or SPG integrity. When we tested miR-285, miRNA known to affect the SPG in the brain lobe, the miR-285 sponge failed to recapitulate the pSJ defects observed in loss of function mutants (Li et al., 2017).

In our sponge screen, the downregulation of 120 miRNAs (with the exception of miR-125) had no effect on peripheral glia morphology. Our analysis with miR-285, known to be crucial for SJ morphology in the CNS (Li et al., 2017), also led to a lack of phenotypes in both the central and peripheral SPG using the miR-285 sponge. While it is possible that miRNAs do not play a role in peripheral glia, there are other possible explanations. For instance, this might happen due to redundancy or compensation of a miRNA loss of function by other miRNAs. There is growing evidence that miRNAs can also bind in a seed-independent manner to their target mRNAs (Chi et al., 2012). Therefore, existing algorithms elucidating mRNA targets of miRNAs based on seed sequence matching might not be able to predict their mRNA targets accurately and subsequently increase the possibility of redundancy. For instance, a genetic miRNA screen in *C.elegans* revealed that miRNAs can redundantly bind to the same target and compensate for the loss of function of a single miRNA (Alvarez-Saavedra & Horvitz, 2010). Alternatively, miRNA-mediated regulatory mechanisms may play a minor role in peripheral glial development in *Drosophila* similar to how the majority of miRNAs individually are not essential for the viability in *C. elegans* (Miska et al., 2007) and *Drosophila melanogaster* (Fulga et al., 2015). To identify and draw more conclusive results for a role for miRNA role in SPG, knockout mutant lines (Chen et al., 2014) could be systematically analyzed for SPG morphology and function in peripheral glia as an alternative to the use of miRNA sponges.

Out of the 120 miRNA sponge lines tested, downregulation of one, miR-125, in SPG resulted in multiple peripheral glia phenotypes. However, the disruptions in SPG morphology observed during the screen were not observed in a miR-125 loss of function mutant or loss of the entire *let-7-c* locus. Overall, this family of miRNAs do not have a function in SPG, and it appears that the miR-125 sponge results were likely false positives. Previous genetic knockdown screens using sponges have also demonstrated false-positive results (Fulga et al., 2015). One possible mechanism might be due to sequestering of unspecific mRNAs leading to off-target effects. This could also explain our observation with the control scrambled sponge having a high phenotype penetrance. Other studies have found non-specific phenotypes with the scrambled sponge, indicating that expression of sponge constructs itself can have off-target effects. (You et al., 2018). Off-target effects might explain the wide range of phenotypes observed with the miR-125 sponge which ranged from loss of SPG morphology to vacuoles and swelling. To check for potential off-target effects, we blasted the miR-125 sponge sequence using a microRNA sponge generator and tester (‘miRNAsong’) (Barta et al., 2016) and found predicted binding to miR-279, miR-3642 and miR-4914. While the expression of the miR-279 sponge did not give a phenotype (Table 1), the other two miRNAs were not tested in our screen. Therefore, it is possible that the phenotypes observed with the miR-125 sponge were due to a range of non-specific targets. Overall, our study suggests that miRNAs tested are not important for SPG morphology or are redundant in nature, and miRNA sponges may not be an effective means of studying miRNA function in the SPG.

## Acknowledgements

This research was supported by a grant from the Canadian Institutes of Health Research (CIHR). We thank Bloomington Drosophila Stock Center for providing the microRNA sponge stock lines. We also wish to thank Dr. David Van Vactor for gifting us the UAS-9a-sponge line and Dr. Nicholas Sokol for gifting us all the *let-7-c* mutants lines.

